# A comprehensive map of the sequence basis of transcription initiation in humans

**DOI:** 10.64898/2026.10.29.762743

**Authors:** Adam Y. He, Claire Tian, Anshul Kundaje

## Abstract

Transcription initiation is a key marker of active enhancers and promoters. Here we describe an atlas of efficient and interpretable deep learning models trained on over 200 transcription initiation profiling experiments representing all major human organs and many cancer types. We show that our models accurately predict initiation patterns from local DNA sequence and decode their sequence determinants across diverse cellular contexts.

## Main

Gene expression is a critical biological process that is dynamically regulated through the complex interplay of environmental stimuli, protein signaling, chromatin state, and regulatory DNA sequence. In mammalian genomes, control over gene expression is chiefly encoded through millions of *cis*-regulatory elements (CREs), which bind transcription factors (TFs) and other chromatin-associated proteins to regulate the expression of nearby genes [1]. Identifying, cataloging, and decoding the mechanistic basis of these elements has spawned multiple consortium-scale projects and remains a central goal in modern genomics [2–4].

Two of the most common and important classes of CREs are promoters, which directly initiate transcription of genes, and enhancers, which can activate transcription from promoters across long genomic distances. Active promoters and enhancers are defined by a number of characteristic biochemical features, including chromatin accessibility, various forms of post-translational histone modification, and transcription initiation [5–7]. In mammalian genomes, both enhancers and promoters are transcribed in sense *and* antisense directions, resulting in characteristic “divergent” transcription initiation patterns [7–11]. Because many noncoding RNAs, including enhancer RNAs (eRNAs) are intrinsically unstable, their transcription initiation profiles can only be reliably captured using assays that profile nascent capped RNAs, such as PRO-cap [12–15].

Over the last decade, advances in deep learning have enabled the development of computational models that accurately predict molecular phenotypes at base resolution from DNA sequence [16–19]. Model interpretation methods have shown that these models learn a variety of regulatory rules, including sequence motifs, motif composites, and spacing and orientation constraints [16–21]. Base-resolution profile models are especially appealing for transcription initiation because the quantitative signal and spatially resolved profile of transcription initiation carry functional information and have distinct sequence determinants [18, 19]. Unfortunately, previous sequence-to-initiation modeling efforts have been limited to a small number of cell lines with published nascent transcription initiation profiles [18, 19] or collapsed initiation profiles across biological contexts [22]. This narrow training scope has limited our ability to interrogate regulatory grammar across the enormous diversity of human cell states. Recently, the ENCODE Project published a large collection of over 200 PRO-cap experiments spanning diverse cell lines and primary tissues representing all major human organs and many cancer types [23]. Here, we describe an atlas of efficient and interpretable sequence-to-transcription initiation models trained on all PRO-cap datasets released to date by the ENCODE Project.

The PRO-cap model atlas consists of 224 single-task BPNet models [16] (each with 7 cross-validation folds) trained to predict PRO-cap profiles from local DNA sequence in each of the PRO-cap experiments published by the ENCODE Project (Fig. 1a). Our models produce highly accurate predictions of transcription initiation strength (“counts”, Fig. 1b) and positioning (“profile”, Fig. 1c), though we note that model performance can degrade significantly with shallow sequencing depth, in experiments without cap-enrichment, or in experiments with other technical issues (Fig. 1d, Supplementary Fig. S1).

**Figure 1.**
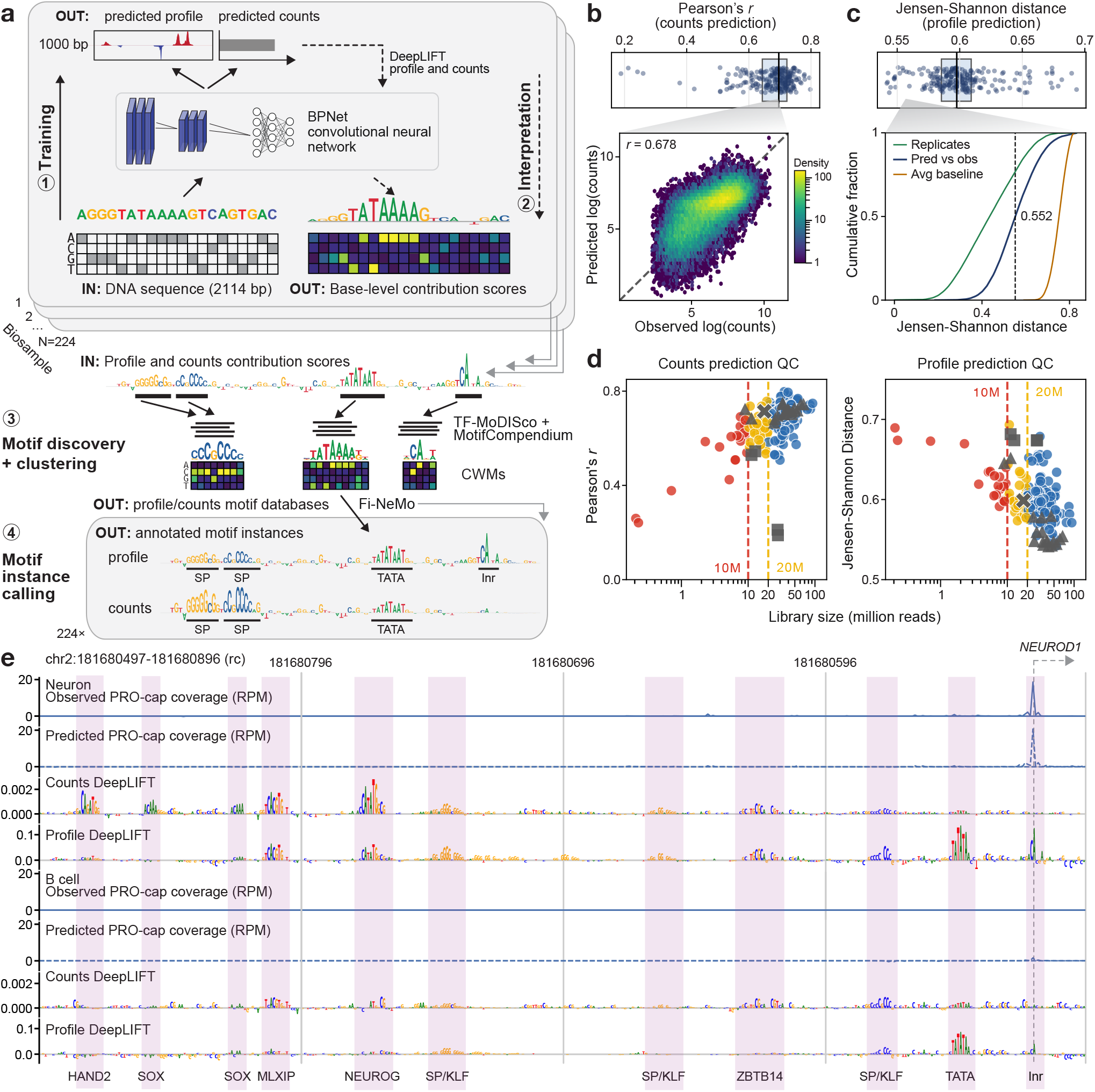
Using deep learning to map the sequence basis of transcription initiation. **a**. Overview of PRO-cap model atlas training and interpretation workflow. **b**. Counts prediction performance. Top, distribution of per-model Pearson correlations between predicted and observed log counts in peak regions per model (mean across seven folds). Bottom, predicted versus observed log counts for neurons. **c**. Top, distribution of median Jensen-Shannon distance between observed and predicted profiles in peak regions per model (mean across seven folds). Bottom, cumulative distribution of neuron profile prediction distances, with replicate and average-profile baselines. **d**. Atlas-wide model quality control and *post-hoc* filtering criteria. Grey markers denote manually flagged or special-case libraries, including perturbation (triangle), uncapped (square), and faulty (cross) samples. All other libraries are colored by read depth (red = fewer than 10 million, yellow = 10 to 20 million, blue = more than 20 million). **e**. Example of the *NEUROD1* promoter in neurons (above) and B cells (below), showing observed and predicted PRO-cap coverage in reads per million (RPM), DeepLIFT scores, and manually annotated motif instances. Counts DeepLIFT scores are rescaled to RPM.

We used DeepLIFT to estimate the contributions of individual nucleotides around PRO-cap peaks to predicted initiation counts and profiles [16, 24]. Visual inspection of these contribution scores revealed a dense landscape of regulatory motifs and divergent regulatory lexicons governing transcription initiation counts and profile (Fig. 1e). Consistent with previous work, we found that transcription initiation profile predictions are largely determined by core promoter motifs such as the TATA box and initiator, whereas counts predictions are driven by sequence-specific TF motifs [18, 19]. We also identified clear instances of cell-type specific TF motifs driving differential transcription initiation strength (Fig. 1e) and positioning (Fig. S2) across cell types.

To systematically identify the sequence drivers of transcription initiation in the human genome, we used TF-MoDISco to cluster high importance subsequences (“seqlets”) into sets of regulatory motifs for each PRO-cap model [25], followed by cross-model meta-clustering with MotifCompendium to obtain an organism-wide, deduplicated lexicon of consensus initiation-driving motifs [26]. We found the profile lexicon to be dominated by core promoter motifs such as the CA and TA initiator motifs, the TATA box, and low complexity motifs likely representing the B-recognition element, the downstream promoter element/motif ten element, or the pause motif, with some contribution from sequence-specific TFs. The initiator and TATA box motifs were highly prevalent and found in nearly all modeled biological contexts (Fig. 2a).

**Figure 2.**
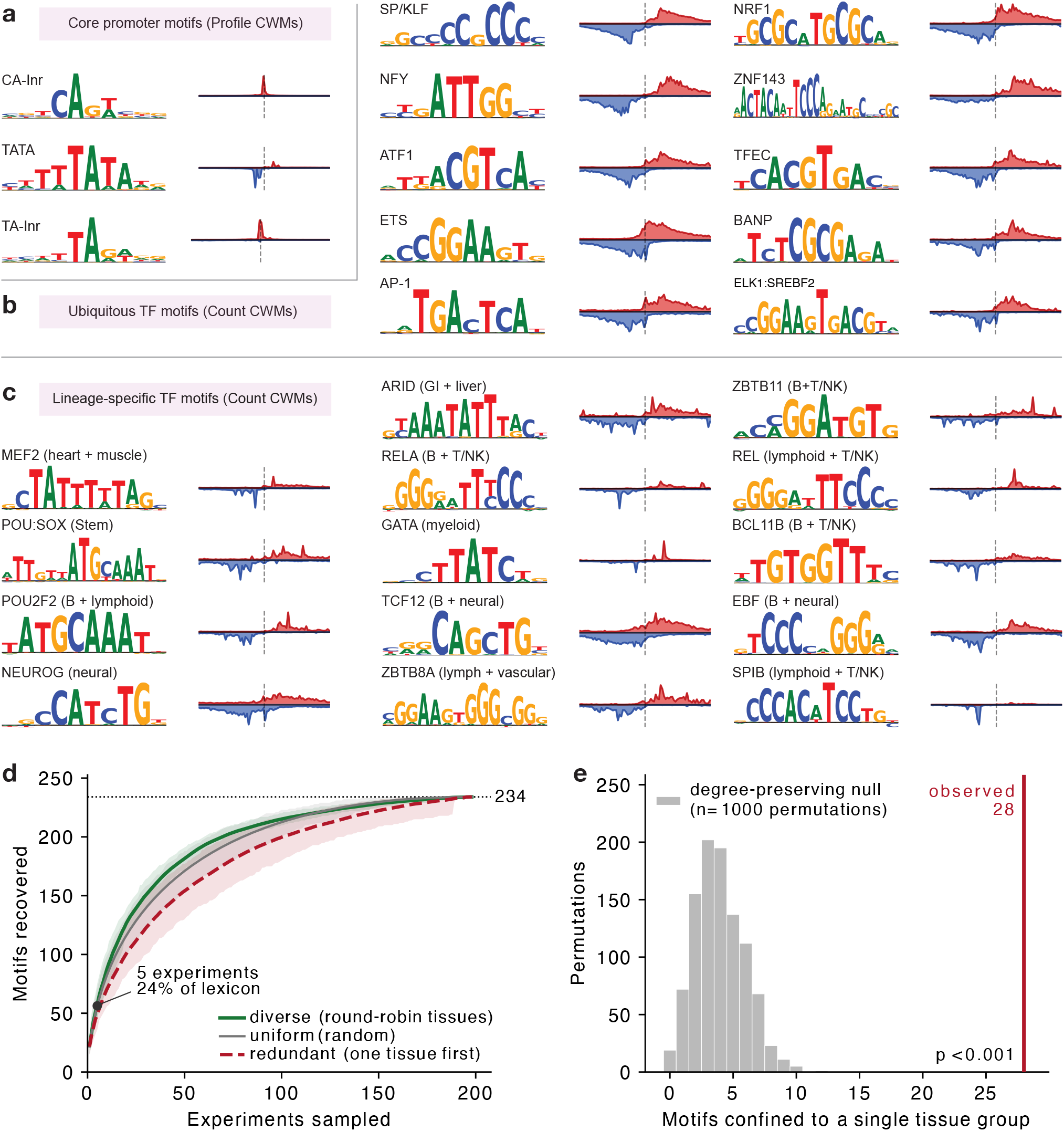
Atlas-scale sequence models recover shared and lineage-specific regulatory motifs. **a–c**. Representative contribution weight matrices (CWMs; sequence logos) from MotifCompendium and motif-centered, strand-specific PRO-cap metaprofiles. CWMs were derived from attribution scores for the predicted initiation profile (**a**) or total initiation count (**b, c**). Shown are core promoter motifs (**a**), broadly recovered TF motifs (**b**) and TF motifs restricted to particular tissue or lineage groups (**c**). Metaprofiles of read density centered seqlets (dashed line, scaled independently per motif) are shown next to the associated CWM. **d**. Motif recovery under tissue-diverse, uniform-random and tissue-redundant experiment sampling; shading shows variation across resamplings and the dotted line marks the complete 234-motif lexicon. Five diverse experiments recovered 24% of the lexicon. **e**. Motifs confined to a single tissue group in the observed atlas (28) compared with 1000 degree-preserving permutations (*p* < 0.001, one-sided permutation test).

We next analyzed motifs discovered by the counts heads. We discovered a diverse lexicon of sequence-specific TF motifs, covering both lineage-specific and ubiquitous TFs (Fig. 2b-c). Lineage-specific motifs recapitulated expected biology, including MEF2 in heart and muscle, GATA in myeloid and erythroid samples, and POU2F2 in B cell and bulk lymphoid samples (Fig. 2c). By contrast, TFs such as the SP/KLF family, NFY, NRF1, and ZNF143 were recovered in nearly all tissue types (Fig. 2b). Motifs exhibited a broad range of positional dependencies (Fig. 2a-c).

Curiously, GATA motifs were absent from the previously described ProCapNet K562 models despite the known critical role of GATA1 in erythroid cell gene regulation [19]. We show that the discrepancy with this work arises primarily because GATA’s role in establishing chromatin accessibility is confounded when both positive and negative training examples are drawn from accessible chromatin, and secondarily because shallow sequencing depth causes disproportionate dropout of the lowly transcribed enhancers that GATA1 predominantly binds [27] (Supplementary Note 1).

After deduplication, motifs detected in at least two experiments formed 234 MotifCompendium clusters (Fig. 2d). In rarefaction analyses, we found that five experiments (similar in scale to previous efforts [18, 19, 22]) recovered only 24% of the cluster-level lexicon. Twenty eight TF-matched clusters were confined to a single tissue group, substantially more than under degree-preserving randomization (7.1 *×*over empirical null, *p* < 0.001; Fig. 2e). Tissue-level concentration persisted after deduplication and reflected lineage restriction across distinct biosamples rather than replicate redundancy (Supplementary Fig. S3a-b).

To determine the extent to which our model atlas captures context-specific sequence effects, we compared predictions within highly tissue-specific peaks (using the *τ* specificity index [28]). Among the top 1% most tissue-specific peaks, predictions ranked the observed dominant tissue group first for 33.4% of peaks (6.7*×* the 5% base rate; Supplementary Fig. S3c). Across pairs of biosamples with replicates, the correlation between predicted and observed initiation differences increased with the replicate-derived reproducibility ceiling (through-origin slope = 0.26; Supplementary Fig. S3d). This suggests that the models encode reproducible context-specific sequence effects but do not recover the full state-dependent range of initiation. We did not observe substantial differences in cross-cell-type initiation prediction accuracy between promoters and enhancers.

Together, these results provide a systematic map of the sequence features associated with transcription initiation across human cellular contexts. The contrast between a broadly shared core promoter grammar that positions initiation and a much larger, lineage-structured TF lexicon associated with initiation strength helps explain how common transcriptional machinery supports context-specific regulatory programs. The incomplete saturation of this lexicon with limited sampling emphasizes the value of expanding initiation profiling across tissues and cell states. The limited agreement between predicted and observed differences across biosamples shows that substantial reproducible variation remains unexplained by these local sequence-only models. By making the models, predictions, nucleotide-resolution contribution scores, motif annotations and genome-browser tracks broadly available, this atlas provides a resource for TSS annotation improvement, noncoding variant scoring, regulatory syntax discovery, and regulatory sequence design.

## Methods

### Data acquisition and preprocessing

We downloaded uniformly processed peak calls (PINTS bed format [15]) and stranded coverage tracks (5’-mapped read counts, bigWig format) from all PRO-cap experiments available on the ENCODE portal in February 2026 (*n* = 224). For experiments with multiple processing releases, only the most recent set of data were retained and any duplicate “archived” files were excluded. Bidirectional and unidirectional peak files were concatenated to produce a single PRO-cap peak set per experiment (bed3 format). One experiment, ENCSR683YIA (Pfeiffer cells), only had “divergent” peak calls available; this was used as the sole source of peaks for that experiment’s models. Signal tracks from technical replicates were summed to produce single positive and negative strand tracks for each experiment. We note that negative strand coverage tracks are represented as negative values by the ENCODE PRO-cap pipelines. We retain this convention for our merged signal and prediction bigWig tracks, but absolute value the observed tracks during model training and evaluation. All datasets were downloaded in hg38/GRCh38 coordinates and the primary ENCODE hg38 fasta file (https://www.encodeproject.org/files/GRCh38_no_alt_analysis_set_GCA_000001405.15/) was used as the source for all genomic sequences.

Peak intervals were represented by their midpoints. We excluded loci that overlapped the ENCODE hg38 blacklist or lacked a complete 2114-bp input window. Within each training split, peaks with total absolute PRO-cap signal exceeding 1.2 times the 99^th^ percentile were also removed. For each experiment, negative regions were sampled from chromosomes 1–22 and X to match the GC-content distribution of its peaks in 2-percentage-point bins. We note that this is distinct from the DNase hypersensitive site negative sets used by ProCapNet [19]; see Supplementary Note 1 for a comparison. Peak-containing windows were excluded, as were candidate negatives whose total absolute PRO-cap signal in the central 1000 bp exceeded one-half of the first percentile of signal across that experiment’s peaks. Candidate negatives containing more than 10% non-canonical bases were discarded during matching, and the stricter input-validity filters above were applied during training. Training batches contained peak and negative examples at a 7:1 ratio.

### Dataset quality control and filtering

We trained, interpreted, and performed motif discovery on models for each of the 224 ENCODE PRO-cap datasets described above, as any dataset-specific quality issues would only affect the corresponding single-task model. For our integrative analyses and to guide external users of our models, we labeled models with audit flags based on quality control metrics and experimental conditions of their associated PRO-cap datasets. First, due to the impact of read depth on model quality and motif discovery (Fig. 1d, [17]), we flagged datasets with < 10M (*n* = 21) and < 20M reads (*n* = 38 between 10 and 20M reads). Next, we flagged datasets corresponding to experiments with perturbations or uncapped enrichment in their metadata. These flags corresponded to a series of experiments in HCT116 (perturb *n* = 8), HEK293T (perturb *n* = 1), and K562 (uncap *n* = 4). Finally, we performed a manual inspection of metaplots around GENCODE (v49) TSSs [29]. The vast majority of PRO-cap datasets exhibit strong sense-strand enrichment exactly on GENCODE TSSs (Supplementary Fig. S1a). Unsurprisingly, uncapped experiments exhibited poor TSS enrichment (Supplementary Fig. S1b), and libraries with low sequencing depth had very noisy profiles (Supplementary Fig. S1c). We also identified one dataset (ReNcell CX, ENCSR973QQI) with anomalous enrichment of PRO-cap signal downstream of GENCODE TSSs (Supplementary Fig. S1d); this last dataset was manually flagged for exclusion from our integrative analyses (exact criteria per analysis described below).

### Model architecture, training, and evaluation

We trained an experiment-specific BPNet model ensemble for each of the 224 PRO-cap experiments using the bpnet-lite package [30]. Each model receives a 2114-bp one-hot-encoded sequence and predicts PRO-cap signal across the central 1,000 bp. The network used 512 convolutional filters and eight dilated residual layers, without control tracks. The profile head produces two 1000-position strand-specific logit vectors, normalized jointly across strands and positions. The count head predicts a single log-transformed total count across both strands. Models were optimized using the sum of the multinomial negative log-likelihood of the observed peak profile and 100 times the mean-squared error of the total log count. Profile loss was evaluated on peak examples, whereas count loss included both peaks and negative regions.

We used seven chromosome-based folds. For each fold, one chromosome set was held out for testing, the next fold was used for validation and the remaining five were used for training. Models were trained for 50 epochs with AdamW, a batch size of 64 and a learning rate of 0.0005 without scheduling or early stopping; the checkpoint with the lowest combined validation loss was retained. Training examples were randomly shifted by up to 200 bp and reverse-complement augmented. Unlike ProCapNet, we did not mask poorly mappable output positions [19].

Performance was evaluated only on chromosomes held out from the corresponding fold. Count performance was measured by Pearson correlation between predicted log counts and log(1 + observed counts), and profile performance by the Jensen–Shannon distance between the joint strand-specific predicted and observed profiles. Statistics were summarized across the seven held-out folds. Unless otherwise stated, downstream predictions and DeepLIFT scores were averaged across all seven models; the cross-cell-type analysis instead used the single model that held out each chromosome. Forward/reverse-complement prediction averaging was not applied.

### DeepLIFT feature attribution

DeepLIFT and the related DeepSHAP are popular feature attribution methods that have been extensively used to interpret sequence models [24, 31, 32]. DeepLIFT/SHAP assigns a numerical contribution score for each feature (base) to a given scalar output of a neural network model by decomposing the difference between the prediction of an input of interest versus that of a neutral reference (or collection thereof) into an additive combination of contribution scores for all features:

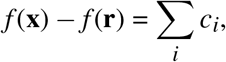

where *c*_*i*_ denotes the contribution of feature *i* in an input **x** to the model output prediction *f* (**x**) compared to model prediction *f* (**r**) based on a reference input **r**.

For explaining DNA sequence models, it is common to use *k*-mer shuffling of the input sequence to generate neutral references [17–19, 24, 31, 32]. Dinucleotide shuffling is particularly popular for regulatory models, as this preserves dinucleotide frequencies (notably CpG) which are correlated with regulatory activity, but destroys more complex TF motifs. However, the effect of reference choice has not been extensively benchmarked, and other references such as mononucleotide shuffles, nucleotide composition, all-zeroes, all-0.25, are conceptually reasonable and have been proposed or used elsewhere [16, 24, 30]. Indeed, there are good *a priori* reasons to suspect that di- or mononucleotide shuffling input sequences is inappropriate for transcription initiation models. Many core promoter motifs are short or degenerate (the main human initiators are CA and TA, and simple AT-rich sequences can drive TBP binding [18]), and so can be inadvertently recreated by shuffling. We show that this causes significant instability in DeepLIFT attributions of PRO-cap models, particularly on the profile head, in Supplementary Note 2.

To bypass this issue, we instead opted to use a deterministic soft reference where each position is set to the nucleotide composition vector of the input sequence [24]. That is, if **x** = (*x*_*b*ℓ_) ∈ {0, 1}^4*×L*^ is a one-hot encoded sequence, then the reference 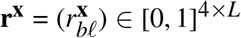 is given entry-wise by

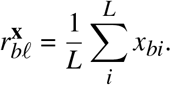

In addition to consistently producing uniform, neutral predicted profiles, a deterministic reference also has the benefit of being much faster to run than random shuffles, which typically must average over some 20 reference sequences to yield stable results even in more favorable settings [17, 32].

Since DeepLIFT can only be computed for a single scalar output, the counts head can be easily attributed, but profile predictions must be aggregated. Following previous work [16–19, 30], we computed profile DeepLIFT scores with respect to the scalar output

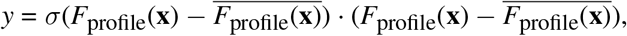

where for a given input sequence **x**, *F*_profile_(**x**) − *F*_profile_(**x**) denotes the zero-centered logits of the profile head (reshaped to a 1D vector), *σ* denotes the softmax operation, and · denotes the dot product.

### Motif discovery, clustering, and annotation

To identify the sequence features driving model predictions, we first computed profile and counts DeepLIFT scores for all models on their respective PRO-cap peaks. We then used tfmodisco-lite [25, 33] to perform *de novo* motif discovery on the DeepLIFT scores from each model and head, resulting in 224 *×* 2 = 448 motif sets. tfmodisco-lite was run over the center 1000 bp of DeepLIFT scores for each peak region, with up to 1 million seqlets per metacluster, and 50 Leiden runs (though we note that 50 is an excessive number carried over from the original ProCapNet and CLIPNET papers [18, 19], and future projects should instead use the standard default of 2 Leiden runs).

Following per-experiment motif discovery, we used MotifCompendium to consolidate profile and counts motifs across experiments [26]. We excluded motif sets discovered from experiments that did not perform cap enrichment (or enriched for uncapped RNAs) or which had anomalous TSS positioning (see previous section on quality control), resulting in 219 motif sets per head. We applied MotifCompendium by first clustering motifs with > 0.95 similarity within each experiment, followed by > 0.9 clustering across all retained experiments. In both steps, we used the Leiden With Constant Potts Quality Function clustering algorithm followed by *k*-centroids refinement. Redundant cluster centroids were iteratively merged by identifying fully connected cliques of centroids exceeding a pairwise similarity threshold of 0.92, followed by k-centroids refinement of individual motif assignments, repeating until memberships stabilized. Resulting MotifCompendium motif metaclusters were matched to known TF motifs by querying against the JASPAR2026 nonredundant vertebrate database [34]. Core promoter motifs, repeats, composite motifs, and other sequences not well-represented in JASPAR were annotated manually.

### Motif hit calling with Fi-NeMo

Motif instances were called separately for the profile and count heads of each experiment using Fi-NeMo [35]. We supplied the full 2114-bp sequence and contribution-score windows centered on each peak and the corresponding experiment-specific TF-MoDISco contribution weight matrices (CWMs). Peak coordinates were matched to the saved sequence and attribution arrays after sequence and blacklist filtering. Fi-NeMo was run with a global regularization parameter of 0.7 and a CWM trimming threshold of 0.3. CWMs trimmed to fewer than six bases were extended symmetrically, within their original boundaries, to a minimum width of six bases. Hits arising from overlapping peak windows were deduplicated before quality control.

We first removed hits in dense clusters of at least five calls of the same motif within 80 bp of a single peak. A preliminary Fi-NeMo report then identified motifs whose hit-derived CWM had similarity ≤0.9 to the discovery CWM. For these motifs, provided they had at least 200 hits, we retained only hits overlapping an independently called seqlet (recursive_seqlets, seqlet-calling threshold 0.05, [32]) in the absolute projected contribution track. The same corroboration requirement was also applied to local profile-head motifs mapped to eight manually identified CA-initiator MotifCompendium clusters, whose short cores could retain high CWM similarity despite excess background hits. After this per-hit filtering, we recalculated CWM similarity and discarded all hits for motifs with a final similarity ≤0.8. Retained hits were then mapped from experiment-specific motifs to their atlas-wide MotifCompendium clusters.

### Motif-centered PRO-cap metaprofiles

Motif-centered PRO-cap metaprofiles in Fig. 2a–c were calculated from TF-MoDISco seqlets rather than Fi-NeMo hits. For each displayed MotifCompendium cluster, we identified the contributing experiment-specific motifs and mapped their seqlets to genomic coordinates using the corresponding peak coordinates and the 1000-bp window used for TF-MoDISco. Seqlets were centered on the informative motif core obtained using the same contribution-weight trimming threshold used for Fi-NeMo hit calling. Where an experiment-specific motif was reverse-complemented relative to the MotifCompendium reference, its seqlet orientations were corrected by aligning its CWM to the cluster reference.

Observed PRO-cap signal was extracted in a ±200-bp window around each seqlet center, oriented to the canonical cluster orientation and averaged in 5-bp bins. Signal was normalized to reads per million (RPM) using the total read count for the corresponding experiment. Seqlets were first averaged within each experiment and the resulting profiles were then averaged equally across contributing experiments, preventing experiments with more seqlets from dominating. The two plotted strands therefore indicate orientation relative to the motif, rather than gene transcriptional sense. Metaprofiles were scaled independently for display, and their amplitudes should not be compared between motifs.

### Motif lexicon rarefaction and discovery concentration

Atlas-level motif analyses were performed on MotifCompendium experiments with at least 10 million mapped reads (*N* = 198 out of 219). For the count head, we retained MotifCompendium clusters represented in at least two experiments. Lexicon recovery was evaluated by sampling *k* experiments without replacement. Specifically, for a motif cluster *c* detected in *n*_*c*_ of *N* = 198 experiments, its probability of being recovered when *k* experiments are sampled without replacement is

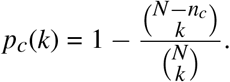

We additionally simulated tissue-diverse and single-tissue sampling schemes, using 200 replicates at each value of *k* and reporting the mean and 5^th^–95^th^ percentile interval. Robustness analyses varied the minimum seqlet abundance used to define motif presence.

For lineage-concentration analyses, the 198 experiments were assigned to 21 tissue or lineage groups. Metastatic samples were grouped by tissue of origin, and blood-derived samples were subdivided into B cell, T/NK cell, bulk-lymphoid and myeloid–erythroid groups to reflect their distinct regulatory programs. We randomized the cluster-by-experiment incidence matrix 1,000 times using the Curveball algorithm [36, 37], preserving both the number of experiments containing each cluster and the number of clusters discovered in each experiment. The observed number of clusters confined to a single group was compared with its permutation distribution using a one-sided empirical *p* value with an add-one correction. The concentration ratio was defined as the observed mean number of tissue groups per cluster divided by its mean across the randomized matrices; values below 1 indicate greater lineage concentration than expected.

### Cross-cell-type prediction analysis

We constructed an atlas-wide peak set by pooling processed PINTS peak intervals from all 224 experiments, retaining canonical chromosomes and merging overlapping or adjacent intervals, yielding 905,540 union regions. For computational tractability, we randomly sampled 100,000 regions using a fixed seed. After excluding blacklists and regions without a valid 2114-bp input window, 99,907 regions remained.

Each of the 99,907 peaks was scored once using the model fold in which its chromosome was held out. Observed and predicted counts were independently rescaled within each experiment to sum to 10^6^ across the analyzed peak set and then transformed using log(1 + *x*). Tissue-specificity analyses additionally excluded peaks whose maximum tissue-group mean was below 0.5 after normalization, leaving 96,121 peaks.

For each peak, let *x*_*g*_ be its mean log-normalized signal in tissue group *g* ∈ {1, … , *G*}, with *G* = 21 as above. We then defined the tissue specificity of each peak using the *τ* specificity index [28]:

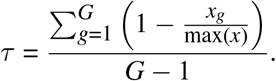

A value of *τ* = 0 indicates equal signal across tissue groups, whereas *τ* = 1 indicates signal restricted to one group. Peaks were ranked by *τ* and divided into the upper 1%, 5%, 10%, 20%, 50% and complete peak sets.

Tissue-retrieval accuracy was calculated as the fraction of peaks for which the observed top-scoring tissue group occurred among the top *k* predicted groups. For each peak, we identified the tissue group with the highest observed mean signal. We then ranked tissue groups by their predicted mean signal and recorded the rank of the observed top group. Top-*k* accuracy was the fraction of peaks for which the observed top group appeared among the *k* highest-ranked predictions. This analysis was restricted to the 20 tissue groups represented by at least two experiments and was repeated across peak sets with increasing tissue specificity.

Differential-signal recovery was evaluated between pairs of biosamples for which both had replicate experiments. For each biosample pair, two independent observed differences across peaks were calculated using disjoint replicate pairs; their Pearson correlation defined the replicate-derived ceiling. The correlation between the corresponding predicted and observed differences was divided by this ceiling to obtain the attained fraction. For comparisons with positive ceilings, we fit the correlation between predicted and observed differences against the replicate-derived ceiling by least squares through the origin, pooling same- and cross-tissue comparisons.

## Data and code availability

The primary code repository for this project is https://github.com/kundajelab/procap-atlas, which includes environment install instructions, data preprocessing pipelines, and model training and analysis scripts/notebooks. Model checkpoints, raw data manifests, processed data, contribution scores, and TF-MoDISco and MotifCompendium motif sets are available on HuggingFace (https://huggingface.co/collections/adamyhe/procap-atlas). PRO-cap coverage and contribution tracks can be visualized on the UCSC genome browser via the track hub https://huggingface.co/datasets/adamyhe/procap-atlas-tracks/resolve/main/ucsc/hub.txt. The raw PRO-cap data for this project are available through the ENCODE project (see the code repository for data manifests).

## Acknowledgments

We would like to thank the labs of Haiyuan Yu and John Lis for generating the PRO-cap experiments as part of the ENCODE Project (particularly Sagar Shah, You Chen, and Aiden Leung for leading the generation and analysis of these datasets); Selin Jessa for allowing us to adapt her model atlas schematic; Valeh Amiri, Kelly Cochran, Salil Deshpande, Ruchir Rastogi, Jacob Schreiber, and Chang Yun for helpful discussions on DeepLIFT, MotifCompendium, and Fi-NeMo; and Maximilian Haeussler and Jonathan Casper for assistance with generating tracks hubs for the UCSC genome browser.

Some of the computing for this project was performed on the Sherlock cluster at Stanford University. We would like to thank Stanford University and the Stanford Research Computing Center for providing computational resources and support that contributed to these research results. The content of this manuscript is solely the responsibility of the authors and does not necessarily represent the official views of their institutions or funding agencies.

## Author contributions

A.Y.H. and A.K. conceived of the project. A.Y.H. designed and implemented the analyses with input from A.K. C.T. performed analyses supporting Supplementary Fig. S2 and Supplementary Note 2. A.Y.H. wrote the main manuscript and figures with revisions from A.K. C.T. wrote Supplementary Note 2 with revisions from A.Y.H. and A.K.

## Generative AI disclosure and reflection

Coding agents (specifically Claude Code running Opus 4.6 and 5 and Sonnet 5 and Codex running GPT-5.5) were used extensively to write the code base underlying this manuscript. We note that the authors A.Y.H. and A.K. have extensive experience in training, evaluating, and interpreting regulatory sequence models (of PRO-cap in particular), and that the generated code largely consisted of piping existing scripts that A.Y.H. had previously written by hand, data scraping, and visualizations based on well-documented existing packages. While the use of the generative AI was key to the completion of this project, domain expertise from the human authors was critical to diagnosing a number of data and model quality issues. We caution readers against uncritical application of generative AI, particularly in more exploratory contexts where there are not good benchmarks or scientific priors against which to sanity check generated text or code.

## Supplementary Figures

**Figure S1:**
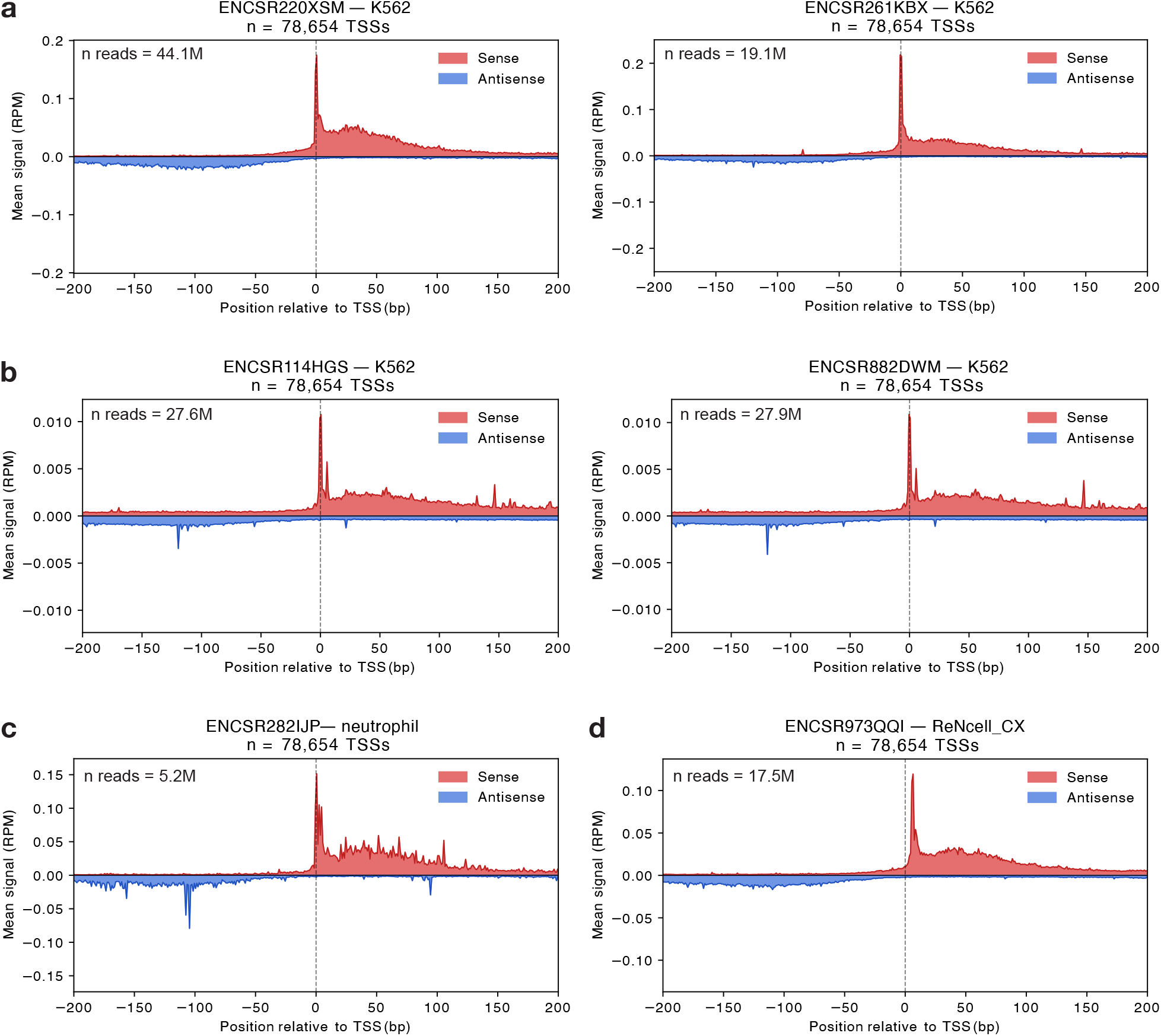
TSS-centered PRO-cap metaplots illustrating dataset quality-control categories. Mean strand-oriented PRO-cap signal across 78,654 GENCODE v49 gene transcription start sites on canonical chromosomes, shown from −200 to +200 bp relative to the annotated TSS and normalized to RPM. **a**. Representative K562 experiments judged good (ENCSR220XSM; 44.1 million reads) or acceptable (ENCSR261KBX; 19.1 million reads). **b**. K562 experiments generated with uncapped-RNA enrichment (ENCSR114HGS and ENCSR882DWM), showing strongly reduced TSS enrichment despite read depths of 27.6 and 27.9 million, respectively. **c**. A very low-depth neutrophil experiment (ENCSR282IJP; 5.2 million reads). **d**. The manually excluded ReNcell CX experiment (ENCSR973QQI; 17.5 million reads), showing anomalous downstream displacement of the sense-strand signal.

**Figure S2:**
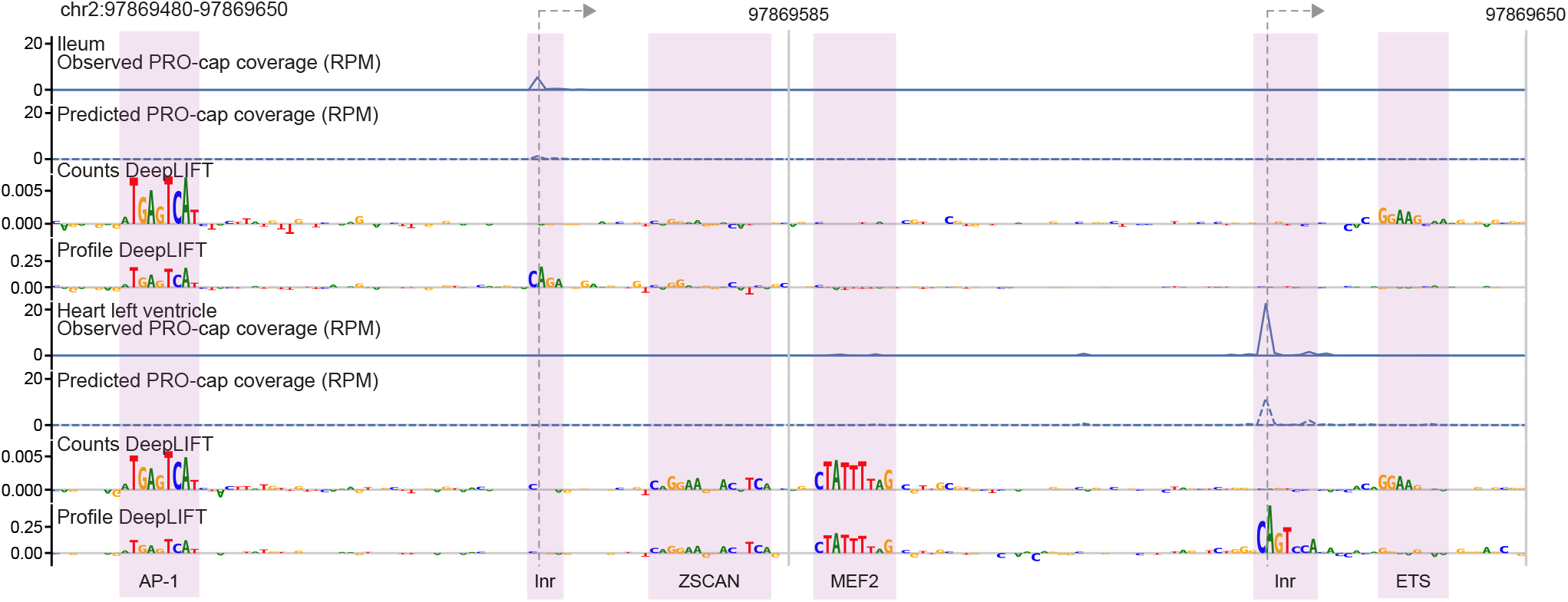
Models implicate cell-type specific TFs in differential TSS usage across tissues. Example of an intronic candidate enhancer (ENCODE SCREEN EH38E2017378 [1], within *TMEM131*) in ileum (above) and heart left ventricle (below), showing observed and predicted PRO-cap coverage in RPM, DeepLIFT scores, and manually annotated motif instances. Counts DeepLIFT scores are rescaled to RPM.

**Figure S3:**
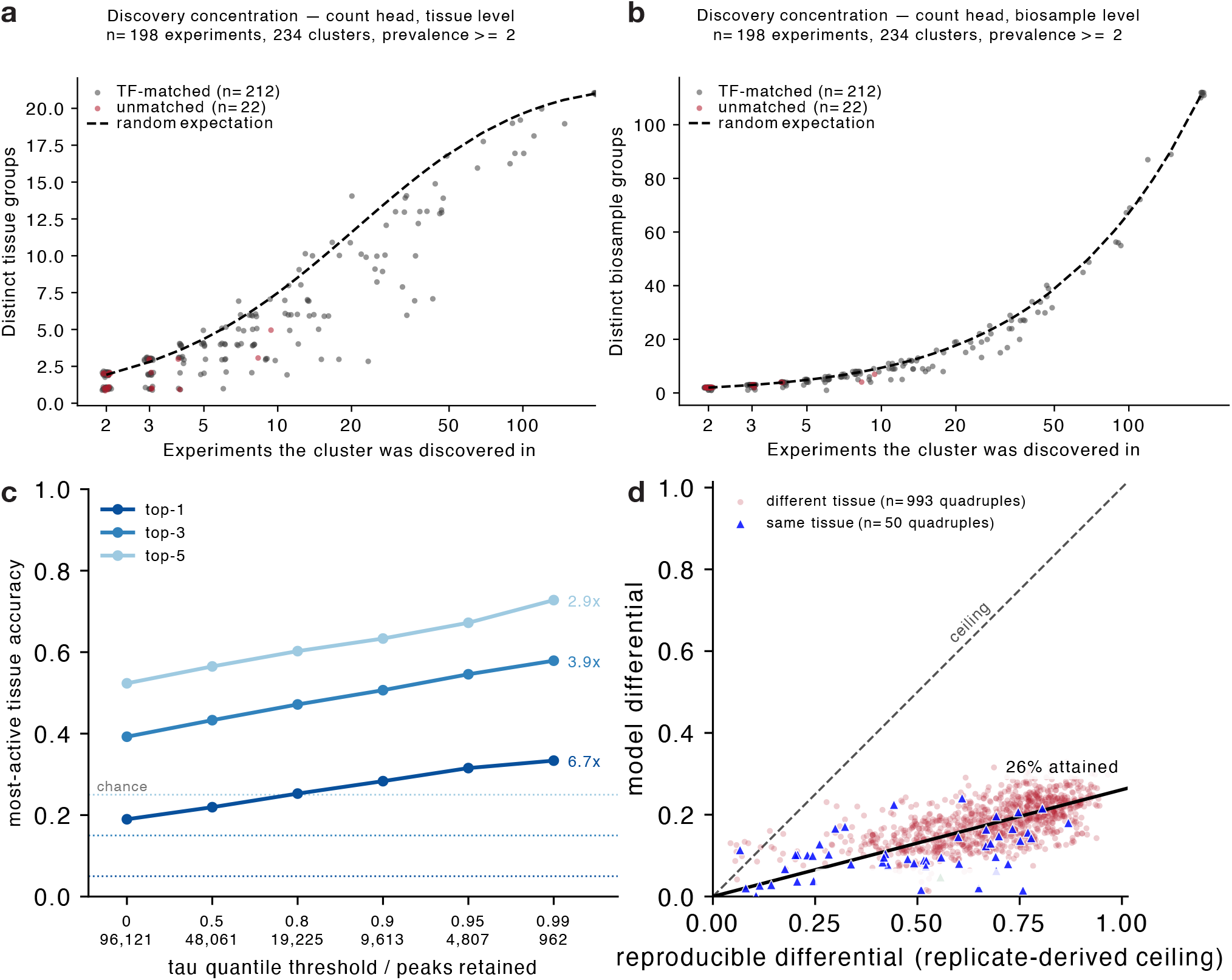
Robustness of motif lexicon estimates and sequence-based prediction of cross-cell type initiation differences. **a**. Discovery concentration on the deduplicated compendium at tissue level (21 groups): distinct tissue groups versus experiment prevalence for each cluster. Points below the dashed expectation curve indicate tissue-concentrated discovery. TF-matched clusters (*n* = 111) and unmatched clusters (*n* = 22) shown separately. **b**. Same as **a** at biosample level (112 groups). Points track the expectation curve more closely, confirming that tissue-level concentration reflects lineage restriction across distinct biosamples rather than replicate redundancy from heavily sequenced cell lines. **c**. Top-1, top-3 and top-5 accuracy for identifying the observed most-active tissue among 20 groups across increasing tissue-specificity *τ* quantile thresholds. Numbers below the axis indicate peaks retained; dotted lines show chance accuracy. **d**. Predicted differential signal plotted against its replicate-derived ceiling for pairs of replicated biosamples. Only positive ceilings are shown; the dashed line denotes the ceiling and the solid line is a through-origin fit (slope = 0.26).

