## Supplementary notes for "A comprehensive map of the sequence basis of transcription initiation in humans"

**Adam Y. He[1], Claire Tian[1], and Anshul Kundaje[1,2]**
Departments of Genetics[1] and Computer Science[2], Stanford University
{ayhe,akundaje}@stanford.edu

### 1 Effect of negative training set and sequencing depth on GATA motif recovery

A previous study modeling K562 PRO-cap data with a similar BPNet architecture noted the absence of GATA family motifs from their K562 models, despite GATA1/2 being master regulators of erythroid gene expression in K562 cells [1]. Cochran et al. acknowledged this absence but did not experimentally test its cause. Our atlas models recover GATA robustly in K562: 7710 count-head seqlets in experiment ENCSR220XSM, 1758 count-head seqlets in experiment ENCSR261KBX, which was used to train K562 ProCapNet (Table 1, Fig 1a). We investigated two hypotheses for this discrepancy using a factorial experiment that varied sequencing depth, peak set, and negative training set composition across five K562 model variants (Table 1).

| Model | Signal | Peaks | Negatives | GATA seqlets |
|---|---|---|---|---|
| ENCSR220XSM | XSM (44M) | XSM (56K) | GC | 7710 |
| ENCSR261KBXtracks_ENCSR220XSMpeaks | KBX (19M) | XSM (56K) | GC | 3692 |
| ENCSR220XSMtracks_ENCSR261KBXpeaks | XSM (44M) | KBX (30K) | GC | 3025 |
| ENCSR261KBX | KBX (19M) | KBX (30K) | GC | 1758 |
| ENCSR261KBX_dnase† | KBX (19M) | KBX (30K) | DHS | 113 |

Table 1: **GATA motif recovery across K562 model variants.** Seqlet counts are from TF-MoDISco applied to count-head DeepLIFT attributions. All GATA patterns matched JASPAR MA0036.4 (GATA2). ENCSR220XSM additionally recovered a minor GATA1::TAL1 pattern (33 seqlets). Cross-experiment models pair one experiment’s merged signal bigWigs with the other’s peak set and use GC-matched negatives. †ENCSR261KBX_dnase most closely approximates the training of K562 ProCapNet from [1].

The first hypothesis concerns the choice of negative training regions. ProCapNet used DNase hypersensitive sites (DHS) lacking PRO-cap signal as negatives, whereas our models use GC-content-matched genomic regions. When the negative training set consists entirely of accessible chromatin (DHS), the model effectively conditions on chromatin accessibility: both positive (PRO-cap peak) and negative (DHS) examples share open chromatin, so GATA’s contribution to chromatin state is confounded with the baseline and cannot be recovered through attribution. In contrast, GC-matched negatives are not enriched for open chromatin, allowing the model to capture GATA’s role in the accessibility difference between PRO-cap peaks and background. Consistent with this hypothesis, switching from GC-matched to DHS negatives while holding all other variables constant reduced GATA seqlet recovery by 94% (1758 to 113 seqlets; Table 1, Fig 1a).

The second hypothesis concerns sequencing depth and its disproportionate effect on enhancer peak recovery. GATA acts predominantly at enhancers [2], which produce low levels of nascent transcription [3]. With

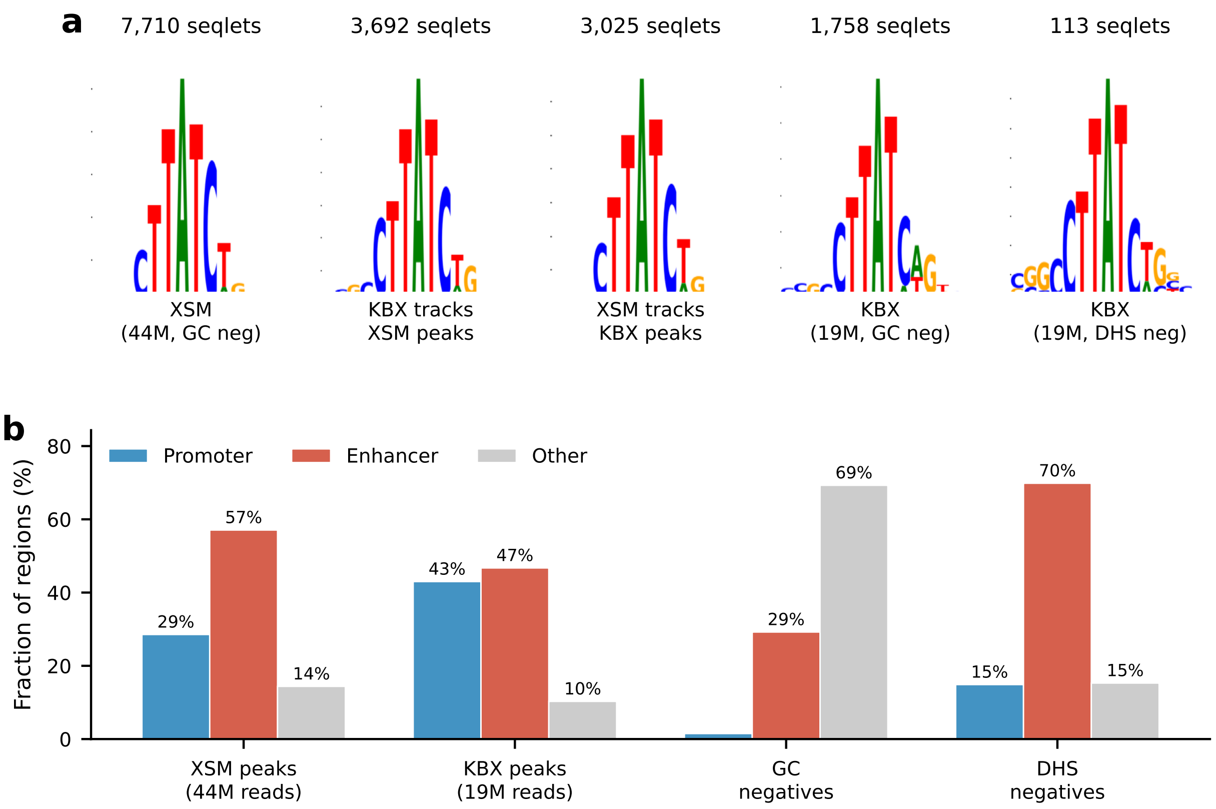


Figure 1: **Effect of negative training set and sequencing depth on GATA motif recovery in K562. a.** GATA contribution weight matrices (CWMs) from TF-MoDISco across K562 model variants, ordered by GATA seqlet count. All variants recover a recognizable GATA motif, but seqlet counts decrease with shallower sequencing, smaller peak sets, and DHS negatives. The DHS-negative model retains only 113 seqlets (vs. 1758 with GC negatives on the same data), consistent with GATA's chromatin-level contribution being confounded with the accessible chromatin background. **b.** Peak and negative region classification against ENCODE SCREEN Registry V4 cCREs. Bars show the percentage of regions overlapping promoter-like (PLS), enhancer-like (pELS/dELS), or other cCRE classes. The deeper ENCSR220XSM experiment recovers proportionally more enhancer peaks than ENCSR261KBX. DHS negatives are heavily enriched for enhancer cCREs compared to GC-matched negatives, as expected for open chromatin regions.

shallower sequencing (ENCSR261KBX, 19.1 million reads vs. ENCSR220XSM, 44.1 million reads), enhancer peaks drop out disproportionately: we observe that the deeper ENCSR220XSM experiment recovers a larger ratio of enhancer peaks relative to promoter peaks compared to ENCSR261KBX (Fig. 1b). This reduces both the number of GATA-containing training examples and the signal quality at the remaining enhancer loci.

To disentangle signal depth from peak set effects, we trained cross-experiment models pairing one experiment's signal tracks with the other's peak set (Table 1, Fig 1a). Training with deeper ENCSR220XSM signal track on the smaller ENCSR261KBX peak set recovered 3025 GATA seqlets (vs. 1758 for the standard ENCSR261KBX model), confirming that improved signal estimates at enhancer peaks boost GATA recovery even without expanding the peak set. Conversely, training with shallower ENCSR261KBX signal on the larger ENCSR220XSM peak set recovered 3692 seqlets, showing that the larger enhancer-enriched peak set also independently improves GATA recovery. Both cross-trained models fall between the two standard models (7710 and 1758), consistent with depth and peak set each contributing partial recovery.

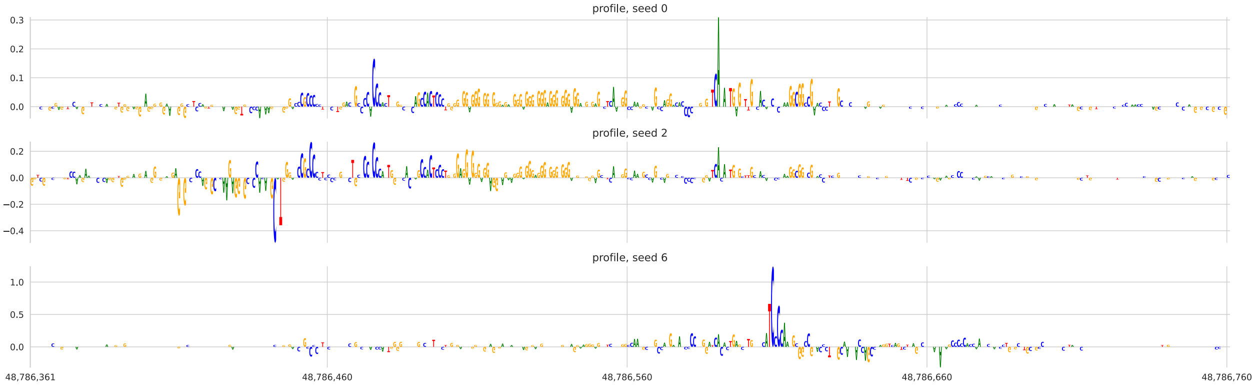

Figure 2: **Sensitivity of profile attributions to dinucleotide-shuffled references at the GATA1 locus in K562.** DeepLIFT/SHAP attribution logos for the same genomic sequence using reference seeds 0, 2, and 6, each generating 20 dinucleotide-shuffled references. Attributions are averaged across the same seven model folds. Changing only the reference seed substantially alters the attribution pattern.

Together, these results demonstrate that GATA motif recovery is sensitive to both the negative training set and sequencing depth, with the dominant effect arising from the choice of negatives. The absence of GATA in previous K562 models should not be used as evidence that GATA does not drive transcription initiation in K562, but rather a consequence of training design decisions that confound GATA's chromatin-level contribution and distal enhancer preference.

### 2 Choice of reference for DeepLIFT/SHAP

DeepLIFT/SHAP quantifies how individual input features contribute to the difference between model predictions for an input sequence and a reference sequence [4–6]. Each feature is assigned a contribution score that reflects how it shifts the model output relative to the reference output, with positive and negative scores corresponding to contributions that increase or decrease the output, respectively. In genomic sequence-to-function models, these contributions are assigned to individual bases in the input sequence. The reference input used for DeepLIFT defines the baseline relative to which nucleotide contributions are assigned. Ideally, an appropriate reference should minimize features relevant to the modeled task and avoid introducing strong model signal of its own. Dinucleotide shuffles of the input sequence are commonly used as references because they preserve mono- and dinucleotide composition, which are important covariates in genomic sequence, while disrupting motifs and higher-order sequence patterns [4–6].

We observed that DeepLIFT profile attributions of PRO-cap models were sensitive to the sampled dinucleotide-shuffled references. At the *GATA1* promoter in K562, changing only the reference seed substantially altered the attribution pattern, even after averaging across twenty dinucleotide shuffles for each of seven model folds (Fig. 2). We hypothesized that this instability arises in part because shuffled references contain core promoter motifs that produce sharp predicted PRO-cap peaks. To test whether these peaks were associated with a shared sequence pattern, we aligned shuffled sequences at the position of maximum predicted PRO-cap signal and computed the nucleotide frequency at each position. This revealed an Inr-like sequence pattern centered near the shuffled prediction maxima (Fig. 3).

To quantify the magnitude of this contamination, we compared the predicted peak strength of 500 dinucleotide-shuffled references against a single frequency reference and the genomic input at five K562 loci spanning a range of expression levels and promoter architectures (Fig. 4): the *GATA1* (chrX:48,786,589), *MYC* (chr8:127,735,434), *ACTB* (chr7:5,530,601; CpG island), and *NFE2* (chr12:54,301,181; no CpG island) promoters and the HS2 enhancer (chr11:5,301,000). Shuffled references produced substantial predicted peaks at all loci, but the severity of the contamination was composition-dependent: the CpG island promoter of

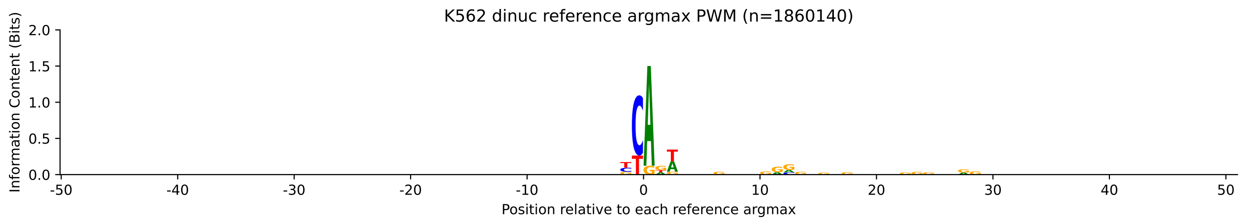


Figure 3: **Position weight matrix at the predicted argmax of dinucleotide-shuffled references in K562.** Shuffled sequences were aligned at their position of maximum predicted PRO-cap signal and nucleotide frequencies were computed at each position (n = 1,860,140 shuffled sequences across all K562 peaks). The Inr-like motif (centered YR dinucleotide) indicates that shuffled references frequently contain core promoter elements that produce sharp predicted peaks.

*ACTB*, which are AT-depleted, exhibited cleaner separation between genomic and shuffled profile prediction, consistent with their depletion of the short AT-rich motifs (such as the Inr) that drive the spurious predictions. At weakly transcribed loci such as the HS2 enhancer, the genomic signal fell within the shuffled distribution, indicating that shuffled references can overwhelm the genomic predictions entirely at weakly transcribed loci.

These results strongly suggest that dinucleotide shuffling, while appropriate for S2F models of epigenomic processes such as chromatin accessibility or TF binding, which are driven by comparatively long motifs (6-20 nucleotides), can fail catastrophically for S2F models of transcription initiation, which is a process that is driven by much shorter motifs (2-4 nucleotides).

A deterministic soft reference, with each position set to the nucleotide composition vector of the input sequence, is a potential candidate replacement for dinucleotide shuffling that has previously been used for attributing S2F models [4]. The frequency reference has a number of drawbacks. Notably, it is out-of-distribution, as S2F are typically only trained on biologically valid, OHE sequences. Moreover, it also fails to account for *k*-mer composition, notably dinucleotide content, which are known covariates for many regulatory sequences. However, for PRO-cap models, it has the very desirable property of preventing degenerate core promoter sequences from contaminating the references.

To quantify attribution stability, we computed pairwise cosine similarities between DeepLIFT attribution vectors generated for the same locus using different reference seeds or model folds. Inter-seed comparisons held the model fold fixed while varying the shuffled reference seed, whereas inter-fold comparisons held the reference procedure fixed while varying the model fold. (Fig. 5). We found that the frequency reference greatly improves inter-fold stability on the profile head and slightly decreases stability on the counts head (Fig. 5).

As a result, we have chosen to implement the frequency reference as the production reference for this manuscript, though we note that other alternatives such as negative sampling should be explored in future studies. We also note that similar problems are likely to arise in the application of DeepLIFT to S2F models other processes such as promoter-proximal pausing or splice site usage that are driven by short sequence motifs, and urge future workers to carefully test for attribution stability.

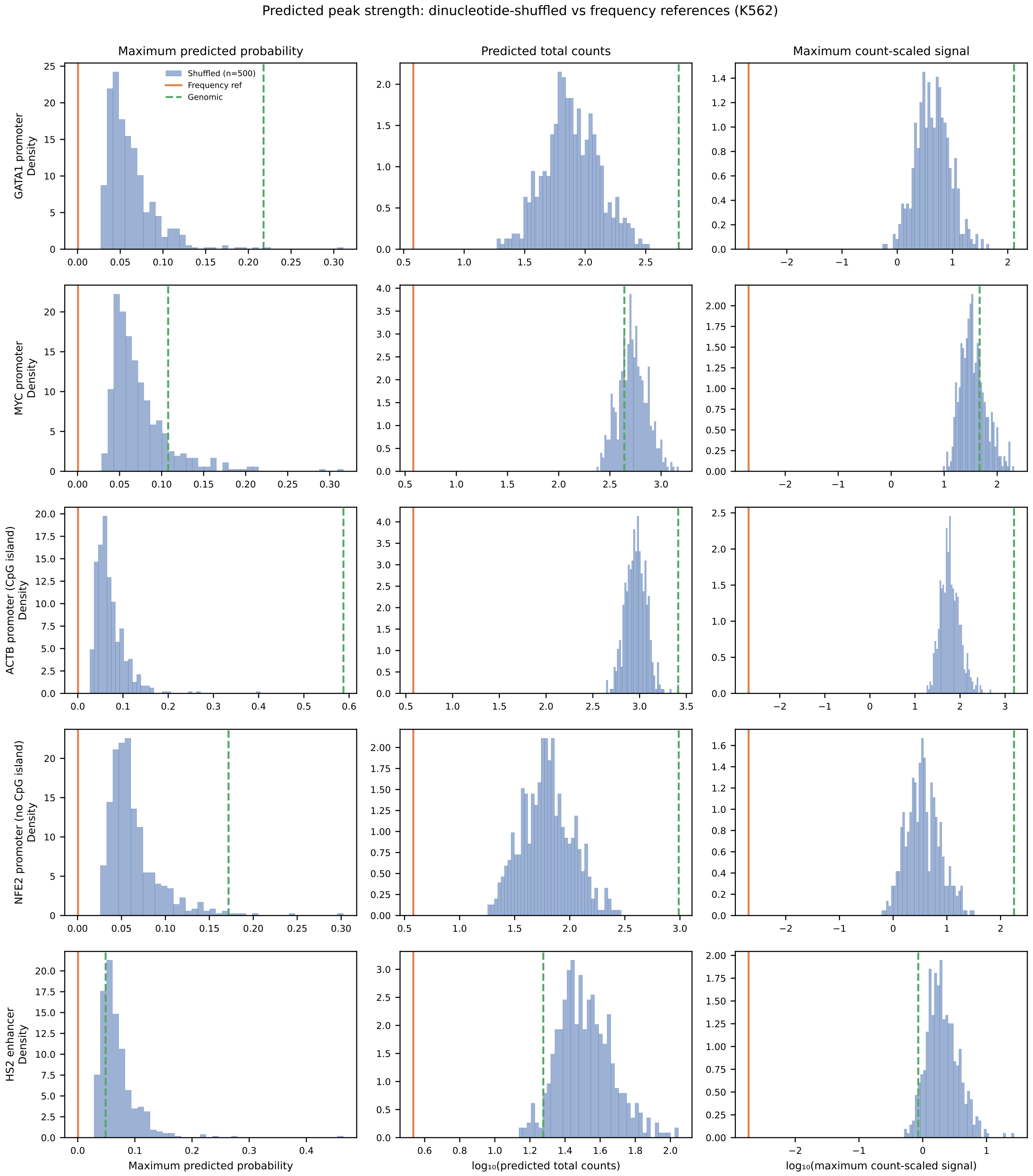


Figure 4: **Predicted peak strength of dinucleotide-shuffled references across five K562 loci.** For each locus, 500 dinucleotide-shuffled sequences (5 seeds $\times$ 100 shuffles) were generated and predicted across seven model folds. Histograms show the distribution of three peak strength metrics averaged across folds: maximum predicted probability (left), predicted total counts (center), and maximum count-scaled signal (right). Vertical lines indicate the frequency reference (orange) and the genomic input (green dashed). Rows from top to bottom: *GATA1* promoter (chrX:48,786,589), *MYC* promoter (chr8:127,735,434), *ACTB* promoter (chr7:5,530,601; CpG island), *NFE2* promoter (chr12:54,301,181; no CpG island), and HS2 enhancer (chr11:5,301,000).

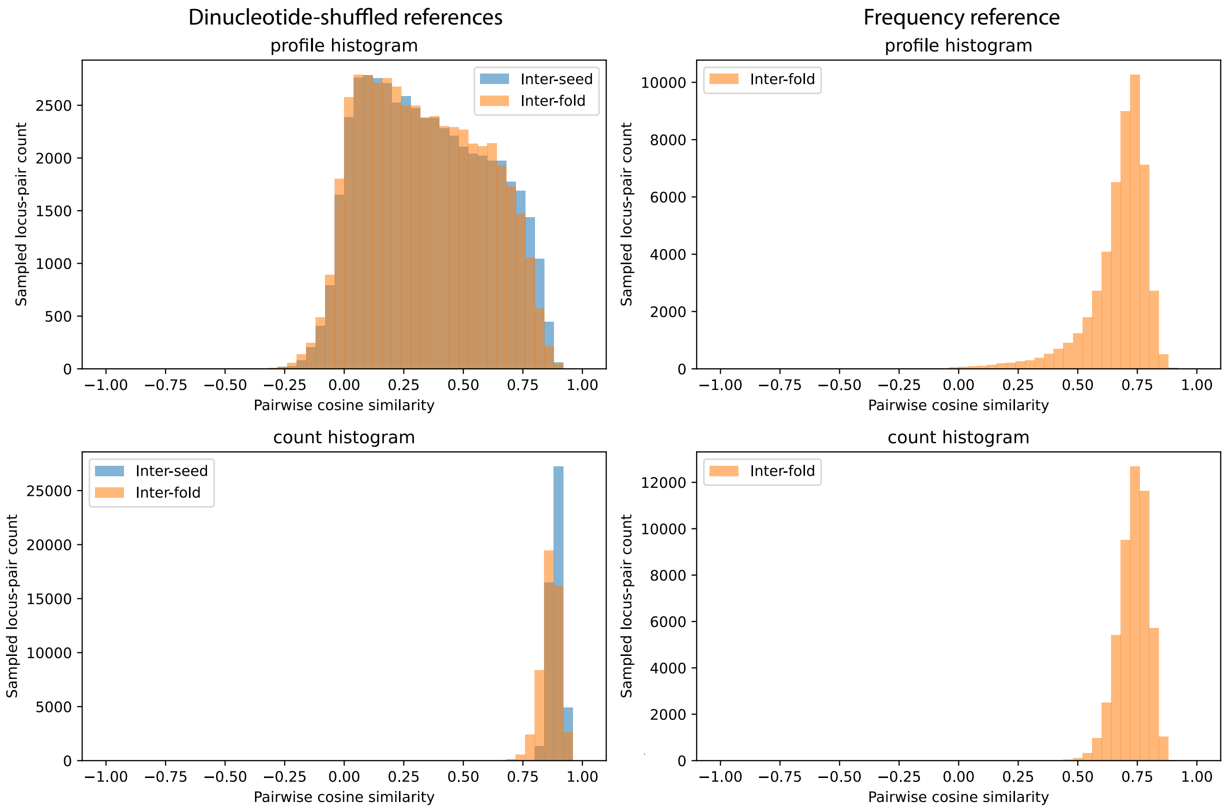


Figure 5: **Stability of DeepLIFT attributions using dinucleotide-shuffled references and a soft frequency reference.** Distribution of cosine similarities between attribution vectors for the same PRO-cap peak across reference seeds (inter-seed) or model folds (inter-fold) in K562. The composition reference greatly improves inter-fold stability on the profile head and slightly decreases stability on the counts head.
